# Rapid disruption of the cortical microcirculation after mild traumatic brain injury

**DOI:** 10.1101/788455

**Authors:** Ellen D. Witkowski, Şefik Evren Erdener, Kıvılcım Kılıç, Sreekanth Kura, Jianbo Tang, Dmitry D. Postnov, Esther Lee, Sunnie Kong, David A. Boas, Ian G. Davison

## Abstract

Traumatic brain injury (TBI) is a major source of cognitive deficits affecting millions annually. The bulk of human injuries are mild, causing little or no macroscopic damage to neural tissue, yet can still lead to long-term neuropathology manifesting months or years later. Although the cellular stressors that ultimately lead to chronic pathology are poorly defined, one notable candidate is metabolic stress due to reduced cerebral blood flow (CBF), which is common to many forms of TBI. Here we used high-resolution *in vivo* intracranial imaging in a rodent injury model to characterize deficits in the cortical microcirculation during both acute and chronic phases after mild TBI. We found that CBF dropped precipitously during immediate post-injury periods, decreasing to less than half of baseline levels within minutes and remaining suppressed for 1.5-2 hours. Repeated time-lapse imaging of the cortical microvasculature revealed further striking flow deficits in the capillary network, where 18% of vessels were completely occluded for extended periods after injury, and an additional >50% showed substantial stoppages. Decreased CBF was paralleled by extensive vasoconstriction that is likely to contribute to loss of flow. Our data indicate a major role for vascular dysfunction in even mild forms of TBI, and suggest that acute post-injury periods may be key therapeutic windows for interventions that restore flow and mitigate metabolic stress.

## Introduction

Traumatic brain injury (TBI) has become a global health epidemic affecting an estimated 69 million people annually (1). While the bulk of TBI cases are mild, they often lead to significant long-term neuropathology and cognitive impairment despite the lack of macroscopic damage to neuronal tissue (2, 3). The continued lack of clinically-proven treatments can be attributed to the diversity and complex time course of injury responses. TBI can drive suppression of neuronal firing, epileptiform activity, or both in succession (4–6). Injury can also trigger cellular processes leading to longer-term damage, including mitochondrial dysfunction, axonal damage from Ca^2+^ influx (7), and accumulation of hyperphosphorylated tau characteristic of chronic traumatic encephalopathy (2, 8) as well as other disease-related proteins such as amyloid-beta and alpha-synuclein (9, 10).

The root causes driving these diverse injury effects have been difficult to establish. While neuronal damage arises in part from direct mechanical trauma, cytoskeletal damage and diffuse axonal injury, and excitotoxicity, it is also linked to disruptions in cerebral blood flow (CBF) (11). Neural tissue imposes high metabolic demands that are met by a densely branched vascular tree, whose small penetrating vessels and fine capillary networks deliver oxygen and glucose in close proximity to virtually all neurons. CBF and vascular tone are continuously modulated by neural activity through a tightly coupled ‘neurovascular unit’ that plays an increasingly recognized role in disease and injury (12, 13). Decreased CBF following TBI may cause metabolic stress that causes or exacerbates neuronal pathophysiology, and a deeper understanding of the vascular effects of injury will be crucial for developing strategies to mitigate its long-term impact.

Postmortem anatomical studies in both animal models and human patients consistently show thinning, shearing, or folding in smaller vessels, as well as microthrombi and microhemorrhages, (14–17). In human TBI, neuronal necrosis correlates strongly with the density of microthrombi (17), suggesting that vascular disruptions impact long-term cellular health and survival. Functional CBF measurements *in vivo* will be crucial for establishing the vascular contributions to mild injury. Reduced flow has been seen in a wide variety of animal models (18–21) and human studies (22, 23), but has largely been examined in more severe forms of injury.

Functional data on mild injuries are limited, particularly during the first hours after TBI that are likely critical in triggering long-term neurodegeneration. In particular, little is known about how mild TBI affects the cerebral microcirculation, the critical site of oxygen and glucose supply (24, 25). Microvascular disruptions can be especially damaging, inducing metabolic stress that quickly interrupts normal neural function and eventually lead to permanent damage or cell death (26–28).

To address how mild TBI alters the cerebral microcirculation, we applied novel label-free imaging methods to visualize flow throughout large volumes of the cortical vasculature *in vivo* at high spatial and temporal resolution. Using repeated imaging with implanted cranial windows, we performed a detailed characterization of CBF changes across both acute (≤3 hours) and chronic (up to 3 weeks) phases of following a closed-skull weight-drop injury. Laser speckle contrast imaging (29) revealed dramatic flow decreases that appeared within seconds to minutes after TBI and persisted for over an hour. Optical coherence tomography (OCT) showed striking dysfunction across the cortical microvasculature, where a large fraction of individual capillaries were occluded for periods of minutes or longer, leading to localized regions with complete loss of perfusion. CBF largely returned to normal by 3 hours after injury and showed no measurable deficits when tracked until 3 weeks. Reduced CBF was paralleled by pronounced vasoconstriction, identifying a potential mechanism contributing to disrupted flow. This rapid and extensive vascular dysfunction highlights the importance of immediate post-injury periods for delivering interventions, and suggests that relieving vasoconstriction will be a promising avenue for countering the effects of mild injury.

## Methods

### Animals and surgical procedures

All experiments were performed in 2-5 month-old male and female C57BL6/J mice in accordance with guidelines of the Boston University Institutional Animal Care and Use Committee. Animals were group housed on a 12-hour light/dark cycle with *ad libitum* access to food and water. Chronic cranial windows were implanted using standard procedures (30, 31). Briefly, animals were anesthetized with isoflurane and a 3 mm craniotomy was made over left primary visual cortex and covered with a glass window composed of 3-4 stacked circular coverslips bonded with optical adhesive (32). The window was sealed with agarose and surgical adhesive (Vetbond, 3M). A stainless steel cylinder (4 mm diameter; 4 mm height) used as target for inducing mild TBI was glued onto bregma (Zap Gel, Pacer). A custom-made stainless steel headpost was attached to the right side of the skull (Loctite 404, Henkel Corp.). The implant was covered with dental cement (C&B Metabond, Parkell). Animals recovered for ≥2 weeks before confirming normal capillary flow, and experimental measurements began at ≥3 weeks post-surgery.

### Mild injury model

Mild TBI was induced using a modified Marmarou model where a weight is dropped onto a metal cylinder attached to the animal’s skull (33) and the animal is unrestrained to allow free movement of the head and body to recapitulate the acceleration and shear forces characteristic of human injury (34, 35). Animals were imaged for ~20 minutes prior to injury, and received buprenorphine analgesia at ~3 minutes before TBI (0.125 mg/kg, subcutaneous). Mice were quickly transferred to a custom TBI apparatus and a 150 g weight was dropped 110-120 cm through a guide tube onto the metal cylinder, propelling the animal through the foil onto foam padding ~7 cm below (5 cm thickness, Mybecca). The weight was secured to fishing line (Stren high impact, Pure Fishing, Inc.) set to a length that prevents potential double-hit injury. Animals were returned to imaging as quickly as possible, within 3-5 minutes of the injury. To facilitate within-animal comparison 5 of 9 mice were given both sham and mild TBI treatments in sequence, where sham animals received identical manipulations apart from weight drop impact.

### Laser speckle contrast imaging

Injury-induced changes in CBF were measured every 1.5 seconds with laser speckle contrast imaging (29). The brain surface was illuminated using a 785 nm stabilized laser diode (LP785-SAV50, 785nm, 50mW, ThorLabs), collimator, and aspheric lens for beam expansion (36). Multiple contrast images were taken at 5ms exposure time with a polarizer and camera lens (VZM 600i, Edmund Optics) and CMOS camera (acA2040-90μmNIR, Basler) and averaged every ~1.5 seconds. Changes in speckle contrast due to local flow-induced scattering were calculated as the ratio of standard deviation of intensity over the mean (29) over a moving 7×7 pixel window using custom software provided by A. Dunn. Speckle contrast images were converted into a blood flow index (BFI) by taking the inverse of the squared speckle contrast, using a custom plug-in for ImageJ (NIH, Bethesda, MD). Relative cerebral blood flow (rCBF) in each animal was quantified for three different regions of interest (ROIs) in the parenchyma spaced throughout the cranial window, where each ROI’s frame-by-frame BFI values were normalized to their mean BFI at baseline (averaged over the 30 frames before injury). The 3 ROIs were then averaged to give a single index of relative CBF for each animal. For further statistical analysis, we compared relative flow values between sham and TBI groups at discrete time points (5, 15, 30, 60, and 120 minutes). 3 animals showing large, rapid (<1-2 sec) fluctuations in speckle contrast due to unstable illumination were excluded from analysis.

### Optical coherence tomography (OCT)

Volumetric imaging of microvascular flow was examined with a spectral-domain OCT system with a 1310 nm center wavelength and 170 nm bandwidth, and a linear CCD (Thorlabs Telesto III) operating at an A-scan rate of 76,000 Hz. With a 10× objective (Mitutoyo Plan Apochromat Objective, 0.28 NA), the axial and transverse resolution was 3.3 μm in the brain. Angiograms were generated with a decorrelation-based method comparing the intensity and phase of two repeated B-scans to identify dynamic tissue(37). Tomographic data was collected with custom LabView routines (National Instruments, Austin, TX). For calculating vessel diameter, angiograms were made from a 1000 ×1000 μm^2^ ROI and scanned with a 1×1 μm^2^ pixel resolution. For examining capillary stalls, angiograms were made from a 600×600 μm^2^ ROI scanned with a 1.5×1.5 μm^2^ pixel resolution. CBF velocity in penetrating vessels was performed by scanning a 1000×1000 μm^2^ ROI with a 2.5×2.5 μm^2^ pixel resolution, using a phase resolved Doppler OCT (prDOCT) which measures the axial blood flow velocity down to capillary level (38) using dynamic light scattering OCT (39), scanning a 1000×1000 μm^2^ ROI with a 2.5×2.5 μm^2^ pixel resolution.

Vessel diameter was quantified using ARIA algorithms in MATLAB (40) applied to angiograms thresholded in ImageJ. A subset of vessels ranging from ~8-60 μm in diameter were measured at all time points, and normalized to their pre-injury size. In Figure 2 D-E and G-H, injury effects were calculated using the average of 5 min and 1 hr time points when changes were maximal). CBF in penetrating vessels was analyzed with custom MATLAB routines (https://github.com/BUNPC/stallingRBCs_OCT) based on manual selection of vessel cross-sections visible at every time point, including thresholding that ensured that results were not biased by ROI size. In 3 TBI animals 72-hour and 1-week time points were not acquired due to hardware errors. The number and duration of capillary stalls were manually identified from angiogram images using custom MATLAB code. Temporarily stalled segments were defined as those alternating between present and absent (i.e., loss of flow signal in >50% of their length) in each ~8.5 minute angiogram time series following previously established protocols (41, 42). Continuously stalled segments were missing during one or more entire time series but reappeared at a later time point.

### 2-photon imaging

Structural stability of the vascular tree was examined using retro-orbital injections of FITC-dextran (50 μl, 5% weight/volume in PBS, Sigma Aldrich) and imaging on a 2-photon microscope (Ultima Investigator, Bruker, Middleton, WI) incorporating a Ti:Sapphire laser (920 nm, Mai Tai DeepSee, SpectraPhysics, Santa Clara, CA) and a 20x objective (Mitutoyo Plan Apochromat, 0.42 NA). Z-stacks of the vasculature were acquired before and after both sham and TBI treatments (~330 μm depth, 3 μm steps). Z-stacks were visualized in ImageJ for manually identifying the same capillary segments observed in OCT images. Subsets of z-stacks containing capillary beds were registered using custom Matlab routines and overlapped images were made in ImageJ.

### Statistical Analysis

All results are reported as mean ± SEM. Statistical significance was calculated using different tests as noted in the results and figure legends. 2-sample t-tests and one-way ANOVAs with Tukey-Kramer post hoc corrections were performed in MATLAB. 2-way mixed ANOVAs with time out to 1 week as the within-subjects variable and condition (sham vs TBI) as the between-subjects variable with Bonferroni confidence interval adjustments were done in SPSS. If a significant interaction between time and condition was detected, a post hoc univariate estimated marginal means analysis was used to determine at which time points sham and TBI differed. p-values below 0.05 were considered statistically significant.

## Results

Although vascular involvement in TBI is well documented, little is known about how injury responses unfold during the first few hours after trauma. The microvasculature, which is responsible for the bulk of neuronal energy supply (24, 25) is particularly understudied. To address how CBF is affected by the mild, closed-skull forms of injury most common in human TBI, we used chronic cranial imaging windows to make longitudinal *in vivo* measurements of the cortical microcirculation, starting within minutes after injury and continuing for up to 3 weeks. Our data reveal striking, rapid-onset deficits in capillary flow that are likely to generate substantial metabolic stress in neuronal tissue, offering a promising target for therapeutic interventions targeting pathophysiology during early phases of TBI.

### Mild injury causes rapid, pronounced decreases in CBF

We first characterized the time course of injury responses by monitoring CBF throughout our 3 mm cortical window with widefield imaging of laser speckle contrast. This approach provides a spatially and temporally precise index of local blood flow averaged across cortical layers based on light scattering from moving red blood cells (29). We acquired an extended image series beginning 20 minutes prior to injury and continuing every ~1.5 seconds for approximately 3.5 hrs.

Mild TBI caused an immediate and dramatic decrease in CBF that was apparent as soon as measurements could be resumed at ~3-5 minutes after injury (Fig 1A). Using the spatial resolution of speckle imaging, we first asked whether CBF was uniformly disrupted throughout the imaging area, consistent with a diffuse mild injury, or alternatively showed any heterogeneity based on local tissue properties. We quantified flow changes for three ROIs chosen to sample widely throughout the imaging window in each animal, centered away from large vessels that can introduce measurement errors. All regions showed virtually identical decreases, indicating that injury causes global reductions in flow across relatively large cortical areas (Fig 1B).

**Figure 1.**
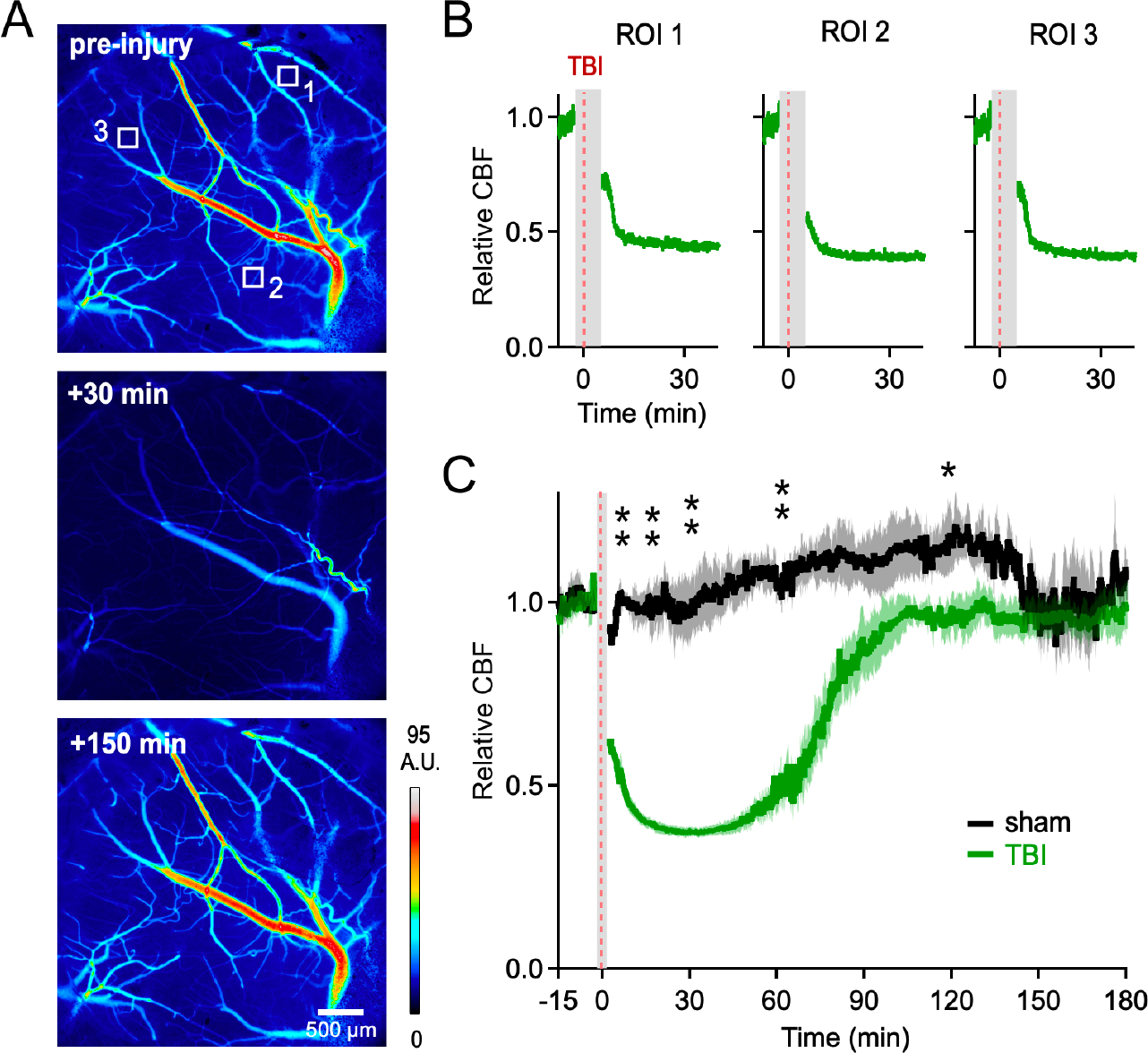
Mild injury causes profound, widespread decreases in CBF. **(A)** Spatial maps of local blood flow index derived from laser speckle contrast imaging before and after injury. Boxes indicate ROIs used to compare flow profile at dispersed spatial locations. Color scale reflects arbitrary contrast units; subsequent analyses are normalized to pre-injury levels. **(B)** Comparing measurements over multiple ROIs showed that decreases in CBF were both extremely rapid and highly similar across locations, indicating uniform effects throughout the imaging area. Red line shows time of injury; gray shading indicates the period when the animal was removed from the imaging apparatus. **(C)** Mean time course of CBF changes pooled across all animals in sham and injury groups, starting 15 minutes before injury and continuing for 3 hours. CBF dropped dramatically within minutes of TBI and remained suppressed for 1.5-2 hours. Solid lines and shading indicate mean ± SEM. ** and * indicate p < 0.005 and 0.05 respectively. n = 4 sham and 6 TBI animals.

**Figure 2.**
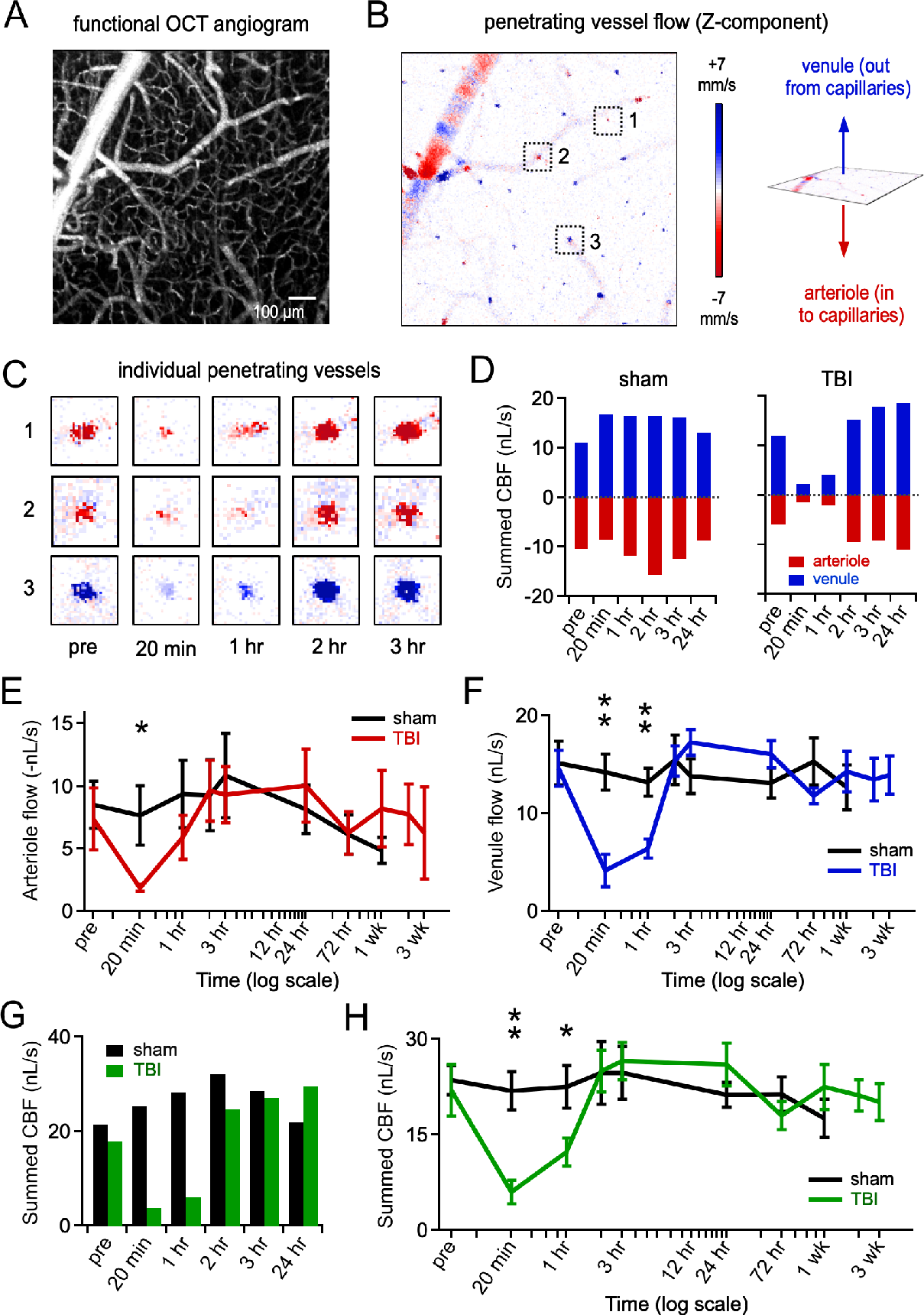
Injury reduces flow in penetrating vessels. **(A)** Angiogram of the cortical vascular tree, shown as a Z-projection of an OCT imaging volume. **(B)** Z-component of flow velocity measured in the same cortical region using dynamic light scattering OCT. Z-velocity is most prominent in vertically oriented penetrating vessels, but also fluctuates at sites adjacent to branches in large surface vessels. **(C)** Examples of flow changes tracked in individual penetrating vessels from the boxed areas in (B), with arterioles and venules distinguished by direction of flow. Images show the first 3 hours; vessels were imaged until 3 weeks post-injury. **(D)** Total flow levels for the full imaging field, calculated by summing the contributions of all individual arterioles and venules. Flow was stable during sham treatment, but strongly suppressed after mild TBI. **(E,F)** Changes in penetrating flow in arterioles and venules respectively, averaged over all animals and all time points. **(G)** Combined arteriole and venule flow for sham and control treatments for the same vessels in an individual animal. **(H)** Total combined penetrating flow averaged across all animals and all time points. ** and * indicate p < 0.005 and 0.05 respectively. n = 7 sham and 7 TBI animals.

To assess the magnitude and timing of CBF changes, we next averaged all ROIs to obtain a single index of flow in each animal, and averaged across all animals in each group. CBF was already substantially decreased as soon as imaging was resumed at 3-5 minutes post-TBI, and ultimately dropped to <40% of pre-injury levels. Flow levels remained depressed by more than half for over an hour before gradually recovering at approximately 1.5-2 hours (Fig. 1C). Quantifying CBF at discrete time points showed a highly significant reduction in TBI relative to sham animals (2-way mixed ANOVA; p < 0.0005 at 5, 15, 30, and 60 minutes; F(1) = 109.6, 430.6, 77.5, and 86.1 respectively; at 120 min: F(1) = 5.56, p = 0.046). In contrast, flow in sham animals remained stable on average (mean across all time points = 104 ± 2% of baseline, one-way repeated measures ANOVA, p = 0.52), and even increased slightly between ~1-2 hours, perhaps reflecting anesthetic effects. Together, speckle imaging revealed a dramatic and widespread decrease in cortical blood supply lasting for over an hour after TBI, suggesting that even mild injury subjects neural tissue to extended metabolic stress through reduced oxygen delivery.

### TBI reduces functional flow in penetrating vessels

Speckle contrast reflects CBF averaged across multiple vessels at different depths and orientations. To directly visualize individual vessels at different levels of the cortical vasculature, we utilized OCT, a label-free imaging approach that provides detailed information about the 3-dimensional distribution of flow based on phase shifts in reflected infrared light caused by red blood cell motion. To specifically quantify how TBI affects flow in the penetrating vessels supplying the cortical capillary network, we turned to phase-resolved Doppler OCT, which measures absolute axial flow velocity in vertically oriented arterioles and venules (Fig. 2A,B; differentiated based on flow direction).

We characterized velocity in ~25 penetrating vessels in each imaging area, and summed their individual contributions (scaled by cross-sectional area) to give an index of total penetrating flow for each animal. Net flow was stable over time in sham animals, but dropped dramatically in both arterioles and venules when assayed at 20 minutes after mild TBI (Fig. 2C,D). Pooled data showed that penetrating flow remained suppressed at 1 hour in both vessel types, with venules still showing a significant reduction, indicating a substantial loss of CBF entering and leaving the capillary bed (Fig 2E,F; arteriole flow for sham and TBI at 20 min = 7.6 ± 2.4 and 1.9 ± 0.3 nL/s; 2-way mixed ANOVA, F(1) = 4.89, p = 0.049; venule flow at 20 min = 14.2 ± 1.8 and 4.2 ± 1.7 nL/s; F(1) = 16.2, p = 0.002; venule flow at 1 hour = 13.1 ± 1.5 and 6.4 ± 1.0 nL/s; F(1) = 14.8, p = 0.002). Flow in both vessel types returned to baseline at 2 hours and remained within a normal range out to 3 weeks. Finally, we tested effects on net flow by combining flow measurements from both arterioles and venules, which showed a similar reduction during the hour after injury (Fig. 2G,H; summed flow for sham and TBI at 20 minutes = 21.8 ± 3.0 and 6.0 ± 1.8 nL/s; 2-way mixed ANOVA, F(1) = 18.494, p = 0.001; at 1 hour = 22.5 ± 3.3 and 12.3 ± 2.2 nL/s; F(1) = 6.422, p = 0.026). Overall, quantitative velocity measurements with phase-resolved Doppler OCT showed strong reductions in the penetrating vessel flow supplying the capillary network that lasted for a period of 1-2 hours.

### Mild TBI causes long-lasting interruptions in capillary flow

Penetrating vessels ultimately supply the cortical capillary network, a crucial vascular element responsible for the majority of O_2_ delivery (24). The size of capillaries is closely matched to red blood cells, which may render them particularly sensitive to flow disruptions. To test how mild injury affects functional flow within the dense capillary network, we used high-resolution OCT angiograms to acquire a time series of 60 imaging volumes, yielding flow measurements every ~7.5 sec for 8.5 min (Fig. 3A).

**Figure 3.**
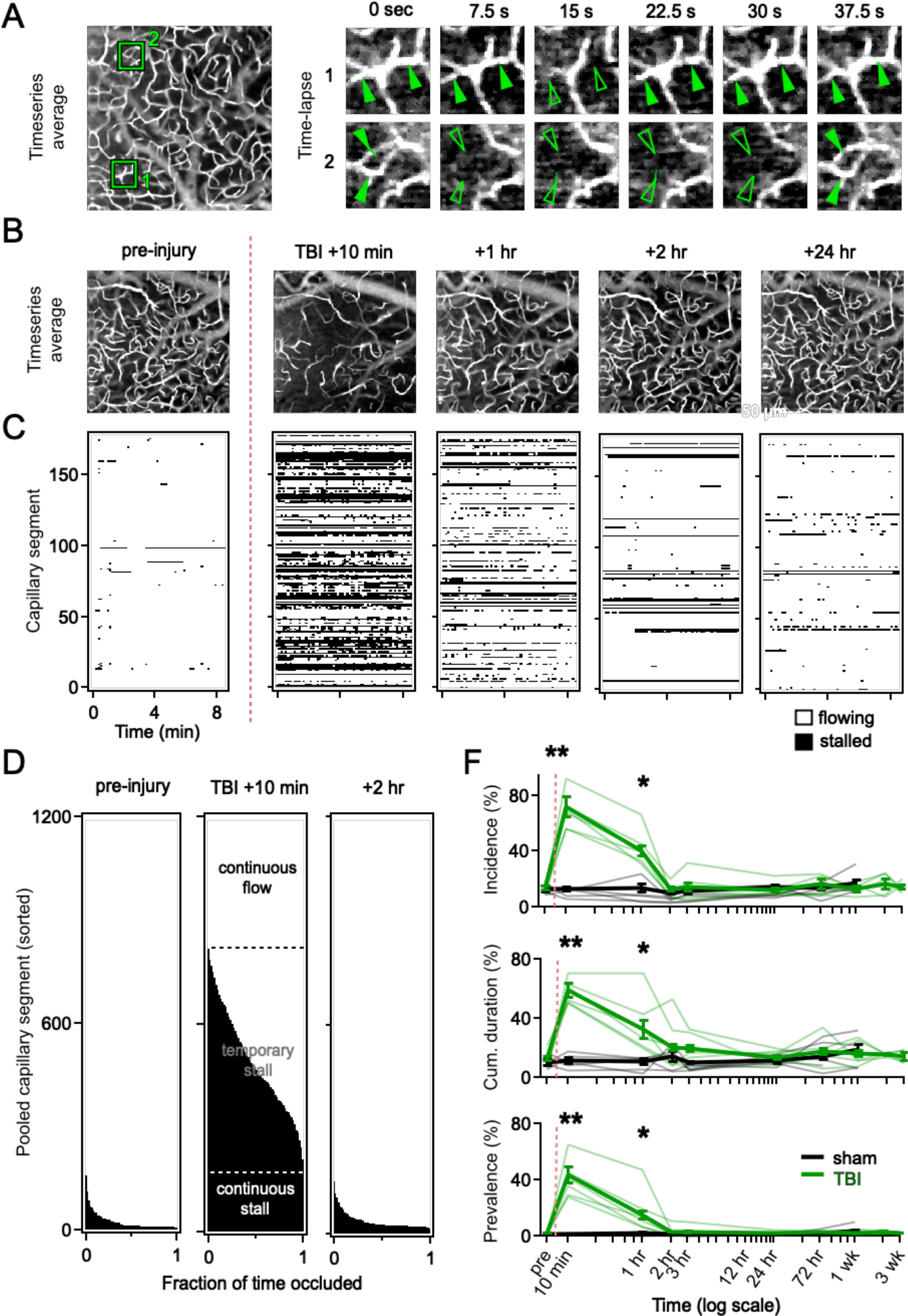
Dramatically increased capillary stalls during immediate post-injury phases. **(A)** Example frames from a time-lapse angiogram sequence showing segments where flow is temporarily stalled (open arrows) and restored (solid arrows). **(B)** Images averaged over all 60 frames of the angiogram time series, shown for a range of initial post-injury time points. Missing segments indicate complete loss of flow during the entire 8.5-minute imaging period. Scale bar = 50 μm. **(C)** Stallogram plots for 181 individual capillaries from the imaging area in (B). Each row shows the behavior of a separate capillary segment over all frames of the time series, with absence of flow indicated in black. **(D)** Cumulative histogram showing the fraction of segments with stalls of varying durations, for all ~1200 capillaries in our dataset. Virtually all segments were continuously flowing prior to injury, while 18% were continuously stalled and 55% only partially functional at 10 min. **(E)** Pooled data showing indices of capillary flow over all time points. Top, percentage of segments showing ≥1 stall during the imaging period (incidence). Middle, percentage of time occupied by flow stoppages in the set of stalled segments (cumulative duration). Bottom, percentage of segments stalled on average at any point during the imaging period. All measures were significantly increased at 10 minutes and 1 hour after mild TBI compared to before injury. ** and * indicate p < 0.005 and 0.05 respectively. n = 7 sham and 7 TBI animals.

Functional angiograms demonstrated robust flow throughout the capillary bed under control conditions. We quantified flow dynamics in capillaries by tracking the presence or absence of OCT signals for 150-200 identified segments in each animal for all 60 frames in the time series. The vast majority of capillaries showed continuous flow throughout, although a small fraction did show brief stoppages lasting several seconds (Fig. 3A), consistent with previous work (41, 42). These ‘stalls’ were infrequent prior to injury, but increased dramatically in both number and duration after mild TBI. In many vessels, we were unable to detect flow at any point in the min imaging series, so that they were entirely absent from the time-averaged images at 10 minutes (Fig. 3B,C). In total we tracked 1189 capillary segments in 7 mice over the course of 3 weeks. At the 10-minute time point, 18% of segments were continuously blocked for the full 8.5 minutes, with an additional ~55% showed intermediate degrees of stalling encompassing a substantial portion of the imaging period (Fig. 3D).

To quantify deficits in capillary dynamics over the full range of post-injury time points, we used several measures. Stall incidence, the percentage of segments showing ≥1 stall at any point, increased from 14% before injury to 72% at 10 minutes and 40% at 1 hour (Fig E; p < 0.0001, repeated measures ANOVA, F(9) = 26.1). In contrast, incidence in sham animals remained low (incidence for sham and TBI at 10 min = 12.8 ± 2.7% and 72.3 ± 5.6%; 2-way mixed ANOVA, F(1) = 92.9, p < 0.0005; incidence at 1 hr = 13.4% ± 4.5 and 40.4% ± 5.0; F(1) = 16.155, p = 0.002). Cumulative duration, the total time any stalled segment remained blocked during the 8.5 min acquisition period, rose from 13% to 59% and 32%, corresponding to 301 sec and 163 sec of stoppage time on average (duration for sham and TBI at 10 min = 11.7 ± 1.6% and 59.2 ± 3.2%, 2-way mixed ANOVA, F(1) = 178.2, p < 0.0005; duration at 1 hr = 11.1 ± 2.0% and 32.4 ± 7.4%, F(1) = 7.698, p = 0.017). Finally, point prevalence, which reflects the average fraction of segments stalled at any point in time, increased dramatically from 2% to 44% and 15% (prevalence for sham and TBI at 10 min = 1.6 % ± 0.4% and 43.7 ± 5.6%, 2-way mixed ANOVA, F(1) = 57.3, p < 0.0005; prevalence at 1 hr = 1.9 ± 0.9% and 15.3 ± 5.7% for sham and TBI, respectively; 2-way mixed ANOVA, F(1) = 5.506, p = 0.037)). The sham group showed no significant changes in incidence, duration, or prevalence. Increases in stalling persisted for over an hour after TBI, but largely recovered by 2-3 hours and remained stable for the ensuing 3 weeks, indicating that effects on the capillary network had a transient time course similar to our other measurements. Together, these data expose a massive decrease perfusion across the cortical microvasculature that is likely to generate extensive, long-lasting interruptions in energy supply to neighboring neurons.

### Mild TBI disrupts vascular tone

While the causes of CBF loss are unclear, one possibility is disruption of vascular tone. Under normal conditions, arterial vessels use smooth muscle to actively regulate their diameter in response to changes in both blood pressure and local neural activity, and pericytes may contribute further constriction at the level of capillaries (43–46). Both dilation and constriction have been described in various TBI models, reflecting disturbances in homeostatic CBF regulation (47–50). However, data examining vascular tone during immediate post-injury periods are limited, particularly for mild closed-skull injuries, and are notably lacking for the penetrating vessels that are the primary site of regulation by neuronal activity levels (45).

Here we characterized changes in vessel size using repeated OCT-based functional angiograms to track the diameter of identified vessels over a wide range of time points beginning minutes before TBI and continuing for 3 weeks post-injury (Fig. 4 A,B). We quantified all vessels in the imaging field that could be repeatedly identified over sessions, spanning a wide range of vessel widths ranging from 8 – 60 μm. Vessels smaller than 8 μm were difficult to quantify and excluded, so that our analysis largely omits capillaries.

**Figure 4.**
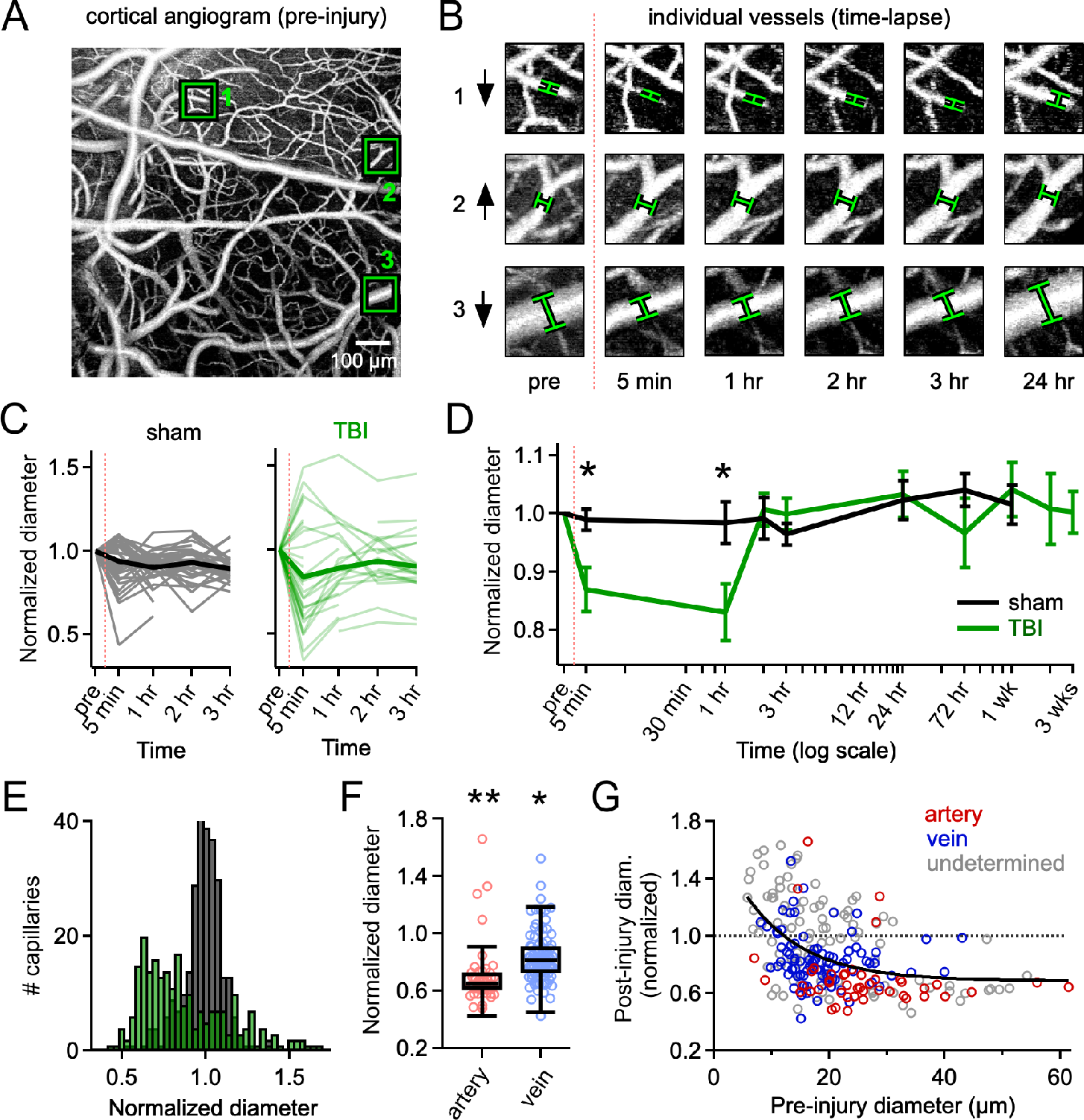
Abnormal vasoconstriction after mild TBI. **(A)** Example OCT angiogram used for measuring vessel diameter. **(B)** Zoomed view of individual vessels corresponding to the boxed areas in (A), imaged over multiple time points to show both constriction and dilation. Upward and downward arrows indicate direction of change during the first hour. **(C)** Normalized diameter for 52 vessels from one animal, shown for both sham and TBI treatments. Injury causes both increases and decreases, but the net effect is constriction. **(D)** Normalized changes in vessel diameter over time, averaged across all animals and all time points, showing significant constriction at 5 min and 1 hr. Points show mean ± SEM. **(E)** Histogram of the distribution of changes in diameter, averaged for both 5 min and 1 hr time points. **(F)** Both arteries and veins showed significant constriction relative to sham, but effects were greater in arteries**. G)** Constriction and dilation depend on initial vessel size before injury, with larger vessels almost always constricting, and smaller vessels showing a mix of constriction and dilation. Black line shows a trendline based on an exponential fit. ** and * indicate p < 0.005 and 0.05 respectively. N = 7 sham and 7 TBI animals.

Interestingly, TBI had non-uniform effects on vessel diameter, which remained stable in sham animals, but showed a mix of both dilation and constriction during the hour after injury, with constriction predominating (Fig. 4B,C). Pooling data across animals, we found that injury significantly reduced mean vessel diameter at both 5 minutes and 1 hour, which we note underestimates the magnitude of injury effects at it reflects partial cancellation of constriction and dilation within each animal (Fig. 4C,D; normalized diameter for sham and TBI at 5 minutes = 0.99 ± 0.02 and 0.70 ± 0.04; 2-way mixed ANOVA, F(1) = 7.10, p = 0.022; diameter at 1 hour = 0.98 ± 0.03 and 0.83 ± 0.05; F(1) = 6.42, p = 0.026). Diameter returned to baseline levels by 2 hours and remained largely stable up to three weeks, paralleling the time course seen with speckle imaging. These findings further suggest that vascular disruptions due to mild TBI occur largely during immediate post-injury phases.

Changes in vascular tone were heterogeneous, biased towards constriction but also including strong dilation (Fig. 4E). This diversity could arise either from variability in injury itself, or from inherent differences in how different vascular elements respond to trauma. While arteries are actively regulated by smooth muscle (51), veins typically respond passively to changes in blood pressure (52). To test for differential injury responses, we identified arterioles and venules based on flow direction in parallel dynamic light scattering OCT data (39). In the vast majority of cases, both vessel types showed clear constriction relative to sham treatments (Fig. 3F: normalized diameter of arteries, veins, and unidentified vessels = 0.70 ± 0.03, 0.84 ± 0.02, and 0.98 ± 0.03, respectively; F(5) = 22.5, p = 2.05e-20, n = 52 and 45 arteries and n = 104 and 87 veins for sham and TBI respectively). Effects were stronger in arteries than veins, consistent with their more robust contractile machinery, and both vessel types had more constriction than their sham counterparts (one-way ANOVA; F(232) = 20.2, p = 8.51e-9). Dilation was largely seen in horizontally oriented vessels where flow direction was not readily categorized.

We further tested whether constriction and dilation may reflect functional subtypes distinguished by size, including larger pial arteries, intermediate arterioles, and small capillaries, where pericytes replace the smooth muscle found in larger vessels (53). Using size as a proxy for vessel type, we plotted normalized post-injury diameter against pre-injury size. Larger vessels ≥30 μm and above showed almost exclusively constriction, while intermediate sizes between 8-30 μm could show either constriction or dilation (Fig. 4H). Dilation largely occurred in the smallest vessels in the distribution, most of which again could not be clearly identified as arteries or veins. Despite relatively high variability, on average we found strong evidence for abnormal, sustained vasoconstriction following mild TBI, identifying a potential driver for the rapid injury-induced loss of CBF.

### Injury has no detectable effects on microvascular anatomy

Disorders such as Alzheimer’s disease and stroke (54) can lead to structural remodeling of the vascular tree that chronically impairs neuronal blood supply (54). Prolonged capillary blockages occur even in healthy tissue, and are linked to subsequent pruning of the stalled segments (41, 42), raising the possibility that the extended stalls we observed could drive similar remodeling of the cortical vasculature.

To test for structural rearrangement, we performed repeated 2-photon imaging of microvascular anatomy labeled with FITC-dextran (55), comparing matched pairs of imaging volumes acquired before and after injury. We first evaluated pruning based solely using the 2-photon dataset, using 3-D image registration to align the pre- and post-injury volumes followed by visual comparison of identified individual segments in the overlaid images (Fig 5A,B). Using this approach, we were unable to detect any evidence of pruning or reorganization at two weeks after mild TBI, with all segments examined being present at both time points.

**Figure 5.**
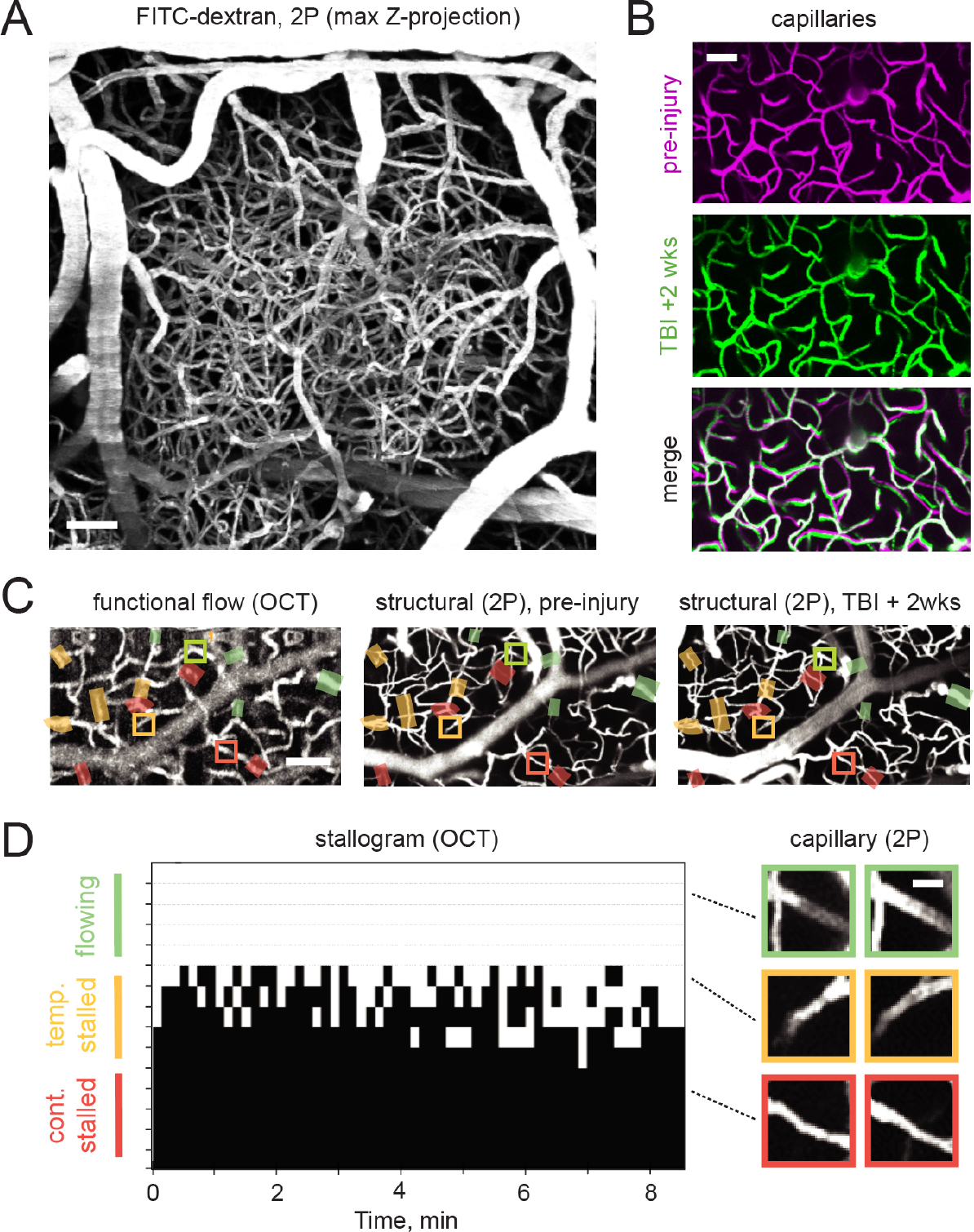
Injury has no discernible effect on structure of the cortical microvasculature. **(A)** Two-photon imaging volume showing vascular anatomy labeled with fluorescein-dextran, shown as a maximum Z-projection. **(B)** Zoomed view comparing the same set of capillary segments before and 2 weeks after TBI. **(C)** Correlated imaging of functional flow (OCT, left) and microvascular structure (two-photon fluorescence, right) in a matched set of capillary segments. Green, yellow, and red shading indicate continuously flowing, intermittently stalled, and persistently stalled segments. Boxes indicate the segments shown in (D). **(D)** Quantification of functional flow stalls (OCT stallograms, left) and structural persistence (two-photon structural images, right). Stalling was quantified for all segments shared between OCT and 2P datasets (~100 per animal); representative examples shown. No capillary pruning was detected in either sham or injury groups, even for segments showing persistent stalls (red).

To further test whether injury may lead to selective removal of a subset of segments with high levels of stalling, which may have been missed by our initial analysis, we correlated our 2-photon data with OCT-based stall measurements for a large set of the same capillary segments (matching OCT data available for an average of 58% of segments, or 99 ± 19 per animal in 5 mice). We classified each of the OCT-matched segments as unaffected, partially stalled, or persistently stalled based on OCT data at the 10-minute time point, and then marked each as present or absent using 2-photon data at the 2-week time point. Again, 100% of the segments we analyzed remained intact, irrespective of the degree of stalling they displayed after injury. Overall, we find that the effects of mild injury are confined to functional flow changes rather than long-term changes in microvascular organization. Together with the near-complete flow recovery we saw by 3 hours, our data imply that any lasting effects of vascular dysfunction arise at the time of injury, rather than from ongoing changes in the microcirculation.

## Discussion

Here we used repeated high-resolution imaging to perform a detailed characterization of injury-induced deficits in the cortical microcirculation *in vivo*, spanning both acute and chronic phases of mild TBI. We describe two features of injury responses that should help inform strategies for restoring flow. First, rapid and dramatic decreases in CBF began within minutes of TBI and lasted over an hour, but showed near-complete recovery by 3 hours. The relatively transient duration of these effects suggest that the timing of interventions may be particularly important in treating mild injury. Second, flow reductions were strongly correlated with changes in vascular tone, including consistently strong constriction in larger vessels. Relieving this constriction may help counter the early effects of injury. Long-term effects on the microvasculature appeared to be minimal, with no evidence of functional flow changes or anatomical reorganization of the microvasculature up to 2-3 weeks after TBI. Together, our results suggest that vascular contributions to long-term pathology are likely to arise at the time of injury via the cellular effects of extended metabolic stress.

### Injury severity and time course of CBF deficits

Vascular disruptions figure prominently in both humans and animal models (23, 56, 57), but their extent and duration can vary widely with injury type and severity. Importantly, CBF changes have largely been examined in open-skull injury models that generate moderate to severe TBI. However, approximately 80% of head trauma cases in humans are classified as mild TBI (58), where vascular effects are likely to differ from those in moderate and severe injury. Furthermore, human data are typically only available several hours after injury. While some animal studies have detected early changes in CBF between 1 hour to 3 days after TBI (18, 19, 21, 59), the full time course of deficits is poorly characterized, and will be important for defining windows for treatment.

Laser speckle contrast imaging allowed us to continuously track flow levels on the timescale of seconds, mapping the full envelope of early CBF disruptions. The onset of deficits occurred within the first minutes or even seconds of trauma, suggesting that rapid interventions may offer potential benefits. CBF continued to decline to levels that were unexpectedly low for our mild injury paradigm, with speckle imaging values falling to ~40% of pre-injury baseline, and OCT-based flow measurements in penetrating vessels reduced to 27% of control. Other mild TBI work has typically found smaller decreases on the order of 60 – 70% of control levels (20, 21, 59, 60). However, most of these studies measured CBF at 4 hours or later (20, 21, 60) and thus may have missed larger initial reductions. While the decreases we found were more severe, they were shorter in duration than seen previously, perhaps due to the relatively long distance between our injury and imaging sites. Injury effects may be even stronger or longer-lasting closer to the site of impact (19). Although briefer, the marked flow reductions we found still lasted well over an hour, making them likely to generate high levels of metabolic stress and physiological dysfunction. Overall, our results suggest that the first 1-2 hours after mild TBI is a key window for vascular effects, a time range that has not been well characterized in mild injury.

### Microvascular disruption on localized scales

Previous studies on CBF and TBI have typically used methods that measure spatially averaged flow. While we also found striking reductions in mean CBF, OCT also allowed us to further visualize flow disruptions in the individual penetrating vessels or capillaries where the vast majority of O_2_ exchange and glucose delivery occurs (59, 61–64). The precise impact of capillary stalls on neural tissue is difficult to predict. Stoppages in the capillary network were all-or-none and highly heterogeneous, and may have even more severe metabolic consequences on local scales. Tissue oxygenation drops off exponentially over distances of tens of microns from the capillaries and pre-capillary arterioles (65, 66), an effect normally countered by dense branching within the capillary network that places most neurons within 15 μm of a vessel (67). However, the widespread and prolonged stoppages caused by injury may substantially increase this distance and create localized pockets of metabolic starvation.

In stroke, occlusion of even a single penetrating arteriole leads to highly localized loss of neuronal function, spreading depolarization, and eventual cell death on scales of ~100 μm (26, 27). Although the neural consequences of finer-scale blockages are unclear, they may be substantial given the 18% of continuously occluded vessels seen 10 minutes after mild TBI. We also found segments that were stalled across multiple time points, suggesting they may have been occluded for 2 hours or more. Additionally, widespread stalls may also prevent the flow redistribution that is normally able to accommodate small focal occlusions (27, 68). Together, our findings suggest that it will be important to consider localized effects when evaluating injury severity.

### Metabolic stress and cellular pathology

A key goal in mild TBI is to identify the biological stressors occurring at the time of injury that ultimately lead to chronic neuropathological changes. Even mild injury can lead to substantial cellular damage and degeneration, especially for repetitive injuries linked to chronic traumatic encephalopathy (8, 69) and Alzheimer’s disease (70, 71). Metabolic stress due to loss of CBF is a prime candidate for triggering cellular pathology. Continuously intact microcirculation is required to meet the high energetic demands of neuronal tissue, and cellular health declines rapidly after CBF is disrupted. In stroke, multiple forms of pathophysiology emerge within 5 minutes of diminished perfusion, including ionic imbalance, structural degradation of the cytoskeleton and mitochondria, and failure of synaptic transmission (72–74). Similar cellular effects are often found in TBI (75, 76), likely due in part from energetic stress from loss of CBF. Neuronal dysfunction and cell death correlate with the degree and duration of CBF loss in ischemia and severe TBI (11, 77). For example, EEG rhythms slow when flow drops to ~50%, with further decreases causing loss of consciousness, followed by disrupted ionic homeostasis, generation of reactive oxygen species by anaerobic metabolism, and eventually permanent neuronal damage (11). Accordingly, extended periods of CBF loss leads to more severe outcomes in human injury (78–81).

Energetic stress may trigger diverse biochemical cascades that continue to cause cellular damage even after flow is restored (7, 76, 82). This includes loss of ionic homeostasis leading to mitochondrial dysfunction (74) and cortical spreading depolarization (83–85). Spreading depolarization increases intracellular calcium, further stressing mitochondria and limiting their ability to generate ATP, buffer intracellular calcium, and negate reactive oxygen species (74), forming a feed-forward loop that worsens oxidative stress. Mitochondrial function can be disturbed for up to a week after TBI (74), leading to accumulation of reactive oxygen species and tau hyperphosphorylation (70, 71).

Overall, decreased CBF is unlikely to fully account for all injury-induced deficits. TBI causes a complex suite of effects, including diffuse axonal injury that degrades both cellular health and neural connectivity (7, 86). However, our findings point to a major role for vascular dysfunction in even mild forms of injury, which has the potential to amplify the mechanical damage arising from initial impact.

### Possible mechanisms and future directions

The primary causes of reduced CBF are unclear. One possibility suggested by our data is pathological constriction of blood vessels, which under normal conditions use smooth muscle to actively regulate their size in response to changes in arterial pressure or metabolic demand (51, 87). TBI caused pronounced reductions of 50% or more in the diameter of both arteries and veins, well below the typical physiological decreases of ~10% (88), and resembling the vasospasm appearing 1-2 days after subarachnoid hemorrhage (89, 90).

Constriction could result from several factors. TBI may act directly on the vascular system itself, disturbing the autoregulatory machinery that normally adapts vessel size to arterial pressure (91–94). In this case, constriction could reflect stretch-induced contractile responses in smooth muscle due to pressure spikes during injury (95, 96), or alternatively hyper-compensation for the later TBI-induced pressure increases seen in fluid percussion models in rats (64, 96). In human TBI, however, both increases and decreases in arterial pressure have been found (97). Alternatively, vessels could be directly compressed by increased intracranial pressure from fast-acting anoxic edema (98).

Finally, extreme constriction could be due to abnormal recruitment of the neurovascular signaling pathways that couple flow levels to local neural activity (31, 99). Pathological bursting, excitotoxicity, or spreading depolarization at the time of trauma may drive abnormal recruitment of constriction (100–102), and there is evidence of this phenomenon in severe human injury (103). Interestingly, constriction due to spreading depolarization has also been seen to coincide with capillary stalling (101, 102)

CBF decreases may also arise at the level of capillaries, whose tight size tolerances may make them particularly sensitive to occlusion during periods of reduced supply from upstream vessels. Capillary stalls occur at low levels even under normal conditions (41, 42), and may be increased by pathological constriction of their surrounding pericytes as seen in stroke and hypoxia (104–106). Inflammatory responses to injury may also increase the “stickiness” of capillary walls, leading to leukocyte and platelet adhesion (17, 47, 107, 108). Blocking inflammatory neutrophil adhesion alleviated both capillary stalls and cognitive dysfunction in a mouse model of Alzheimer’s (109). While it is unclear whether inflammation is rapid enough to account for capillary stalling at 10 minutes, it could easily contribute to later reductions. Capillary disruptions could also contribute to constriction in larger upstream vessels, which may be a normal physiological response that balances stall-dependent increases in intraluminal pressure, or slows blood flow in order to increase oxygen extraction (110).

It will be important for future work to address the biological sources of impaired CBF. Both smooth muscle and pericyte activity can be visualized *in vivo* (111), allowing testing of the role of active contractile processes, or of capillaries vs. penetrating vessels. While technically challenging, CBF measurements could also be combined with imaging of neural activity (112) or local tissue oxygenation (113), which would provide further insight into the metabolic consequences of long-lasting capillary stalls.

Together, our data show that even mild TBI dramatically reduces cerebral blood flow to levels that may subject neuronal tissue to extended metabolic stress, suggesting that the vascular system offers a promising target for mitigating the effects of injury. Flow may be at least partially restored by countering vasoconstriction with vasodilators (89), or by antibody treatments that reduce leukocyte adhesion (109). After recovery of flow, secondary treatments focusing on neuronal pathophysiology, inflammatory responses, and potential reperfusion injury may be more effective, as deployed in stroke (114). While corresponding measurements of CBF in mild human injury will be critical, our data identify immediate post-injury periods as a key window for countering the cellular consequences of metabolic stress.

